# Four high-quality draft genome assemblies of the marine heterotrophic nanoflagellate *Cafeteria roenbergensis*

**DOI:** 10.1101/751586

**Authors:** Thomas Hackl, Roman Martin, Karina Barenhoff, Sarah Duponchel, Dominik Heider, Matthias G. Fischer

## Abstract

The heterotrophic stramenopile *Cafeteria roenbergensis* is a globally distributed marine bacterivorous protist. This unicellular flagellate is host to the giant DNA virus CroV and the virophage mavirus. We sequenced the genomes of four cultured *C. roenbergensis* strains and generated 23.53 Gb of Illumina MiSeq data (99-282 × coverage per strain) and 5.09 Gb of PacBio RSII data (13-54 × coverage). Using the Canu assembler and customized curation procedures, we obtained high-quality draft genome assemblies with a total length of 34-36 Mbp per strain and contig N50 lengths of 148 kbp to 464 kbp. The *C. roenbergensis* genome has a GC content of ~70%, a repeat content of ~28%, and is predicted to contain approximately 7857-8483 protein-coding genes based on a combination of de novo, homology-based and transcriptome-supported annotation. These first high-quality genome assemblies of a Bicosoecid fill an important gap in sequenced Stramenopile representatives and enable a more detailed evolutionary analysis of heterotrophic protists.

## Background & Summary

The diversity of eukaryotes lies largely among its unicellular members, the protists. Yet, genomic exploration of eukaryotic microbes lags behind that of animals, plants, and fungi^1^. One of these neglected groups is the Bicosoecida within the Stramenopiles, which contains perhaps the most common marine heterotrophic flagellate, *Cafeteria roenbergensis^2–6^*. The aloricate biflagellated cells lack plastids and feed on bacteria and viruses by phagocytosis^2^. *C. roenbergensis* reproduces by binary fission and has no known sexual cycle. To our knowledge, the only other Bicosoecid with a sequenced genome is *Halocafeteria seosinensis^7^*, and the most closely related sequenced organisms from other groups are found among Oomycetes and Diatoms. Its phylogenetic position at the base of Stramenopiles and the paucity of genomic data among unicellular heterotrophic grazers make *C. roenbergensis* an interesting object for genomic studies.

Heterotrophic nanoflagellates of the *Cafeteria* genus are subject to infection by various viruses, including the lytic giant Cafeteria roenbergensis virus (CroV, family *Mimiviridae*) and its associated virophage mavirus (family *Lavidaviridae*)^8–10^. We recently showed that mavirus can exist as an integrated provirophage in *C. roenbergensis* and provide resistance against CroV infection on a host-population level^11^. Genomic studies of *Cafeteria* may, thus, reveal new insight into the importance of endogenous viral elements for the evolution and ecology of this group.

Here we present whole-genome shotgun sequencing data and high-quality assemblies of four cultured clonal strains of *C.roenbergensis*: E4-10P, BVI, Cflag and RCC970-E3. The strains were individually isolated from four different habitats (Fig. 1a). Cflag and BVI were obtained from coastal waters of the Atlantic Ocean at Woods Hole, MA, USA (1986) and the British Virgin Islands (2012). E4-10P was collected from Pacific coastal waters at Yaquina Bay, Oregon, USA (1989). RCC970-E3 was obtained from open ocean waters of the South Pacific, collected about 2200 km off the coast of Chile during the BIOSOPE cruise^12^ (2004) (Table 1). In addition, we also sequenced strain E4-10M1, an isogenic variant of E4-10P carrying additional integrated mavirus genomes previously described by Fischer and Hackl^11^. E4-10M1 read data was used to support the E4-10P genome assembly after mavirus-containing data was removed.

**Table 1.** Strain and sample information.

| Species | Strain | Location | Coordinates | Year | Biosample | Roscoff ID |
| --- | --- | --- | --- | --- | --- | --- |
| *C. roenbergensis* | E4-10P | North Pacific, 5 km west of Yaquina Bay, Oregon | 44.62 N 124.06 W | 1989 | SAMN12216681 | RCC:4624 |
| *C. roenbergensis* | E4-10M1 | North Pacific, 5 km west of Yaquina Bay, Oregon | 44.62 N 124.06 W | 1989 | SAMN12216695 | RCC:4625 |
| *C. roenbergensis* | BVI | The British Virgin Islands | 18.42 N 64.61 W | 2012 | SAMN12216698 |  |
| *C. roenbergensis* | Cflag | North Atlantic Ocean, Woods Hole, MA | 41.52 N 70.67 W | 1986 | SAMN12216699 |  |
| *C. roenbergensis* | RCC970-E3 | South Pacific Ocean, 2200 km off the coast of Chile | 30.78 S 95.43 W | 2004 | SAMN12216700 | RCC:4623 |

**Figure 1.**
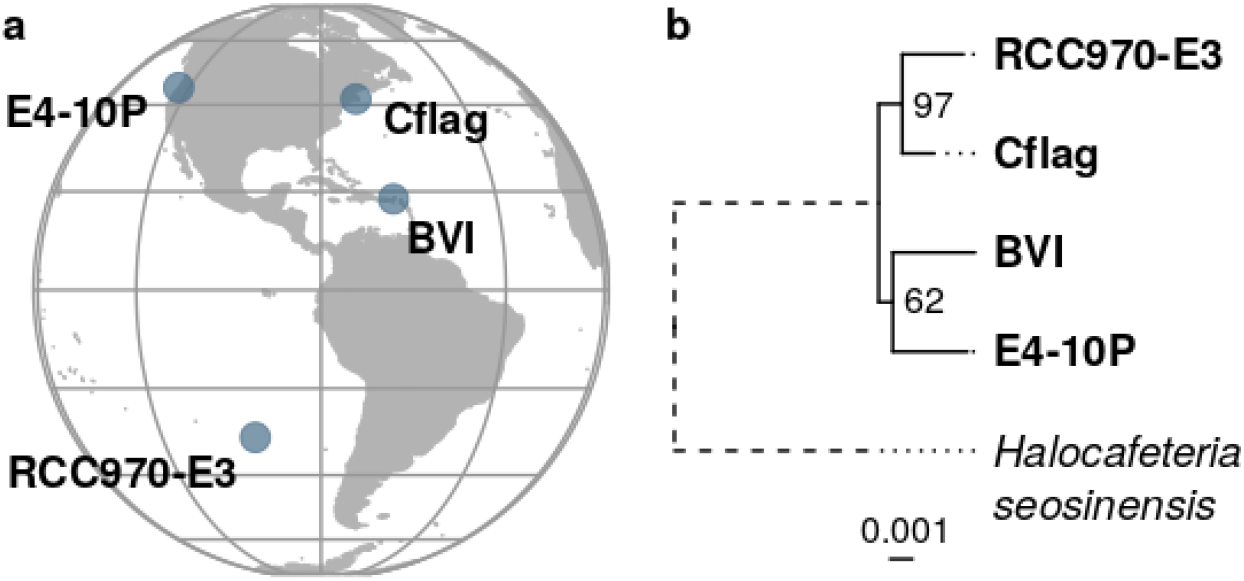
Sampling locations and phylogenetic relationship of *Cafeteria roenbergensis* strains. (**a**) Map representing the sampling sites of the four *C.roenbergensis* strains around the Americas. (**b**) Maximum likelihood tree reconstructed from a concatenated alignment of 123 shared single-copy core genes for the four *C.roenbergensis* strains and their outgroup *Halocafeteria seosinensis.* Numbers next to internal nodes indicate bootstrap support based on 100 iterations. The branch to the outgroup represented by a dashed line has been shortened for visualization.

Overall we generated 23.5 Gbp of raw short read data on an Illumina MiSeq platform with 99-282 × coverage per strain, and 5.1 Gbp of raw long read data on a PacBio RS II platform with 13-45 × coverage per strain (Table 2) (Data Citation 1). Based on 19-mers frequencies, we estimate a haploid genome size for *C. roenbergensis* of approximately 40 Mbp (Fig. 2). We first generated various draft assemblies with different assembly strategies and picked the best drafts for further refinement (see Technical Validation) (Data Citation 3). After decontamination, assembly curation and polishing, we obtained four improved high-quality draft assemblies with 34-36 Mbp in size and contig N50s of 148-460 kbp (Table 3) (Data Citation 2). The genomes have a GC-content of 70-71%, and 28% of the overall sequences were marked as repetitive.

**Table 2.** Sequencing information and library statistics.

| Strain | Instrument | Library layout | # Libraries | Library size (Gbp) | Coverage | SRA study accession | SRA run accession |
| --- | --- | --- | --- | --- | --- | --- | --- |
| E4-10P | Illumina MiSeq | paired | 2 | 6.85 | 171 | SRP215872 | SRR9724619 |
| E4-10P | PacBio RS II | single | 2 | 0.52 | 13 | SRP215872 | SRR9724618 |
| E4-10M1 | Illumina MiSeq | paired | 2 | 4.45 | 111 | SRP215872 | SRR9724621 |
| E4-10M1 | PacBio RS II | single | 2 | 1.3 | 32 | SRP215872 | SRR9724620 |
| BVI | Illumina MiSeq | paired | 1 | 4.31 | 108 | SRP215872 | SRR9724615 |
| BVI | PacBio RS II | single | 3 | 1.8 | 45 | SRP215872 | SRR9724614 |
| Cflag | Illumina MiSeq | paired | 1 | 3.94 | 99 | SRP215872 | SRR9724617 |
| Cflag | PacBio RS II | single | 2 | 0.95 | 24 | SRP215872 | SRR9724616 |
| RCC970-E3 | Illumina MiSeq | paired | 1 | 3.98 | 100 | SRP215872 | SRR9724623 |
| RCC970-E3 | PacBio RS II | single | 2 | 0.52 | 13 | SRP215872 | SRR9724622 |

**Table 3.**
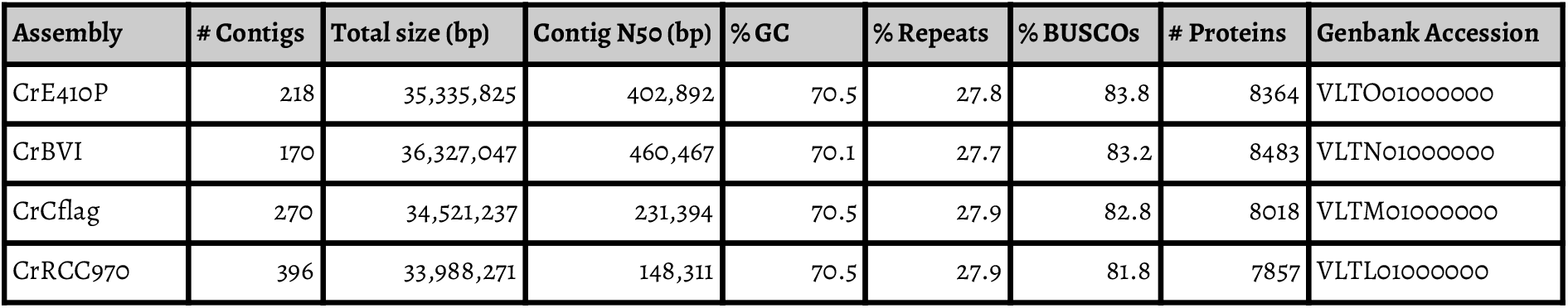
Assembly and annotation statistics.

**Figure 2.**
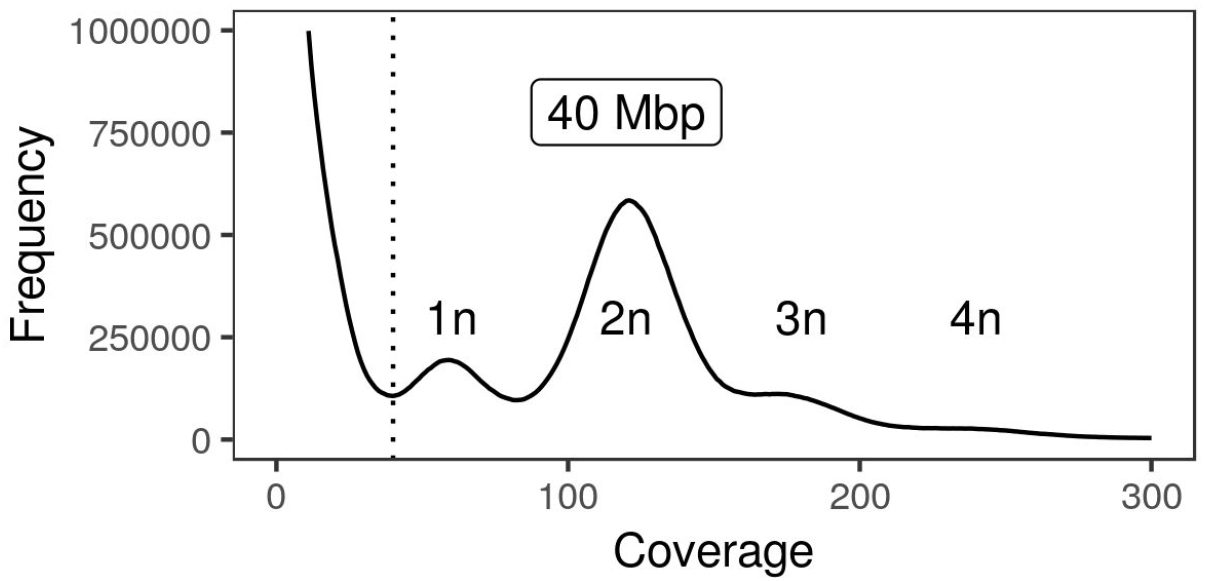
*K*-mer frequency distribution and estimated genome size of *Cafeteria roenbergensis* E4-10P. Frequency distribution of 19-mers in the quality-trimmed MiSeq read set of *C. roenbergensis* strain E4-10P. The major peak at ~120 × coverage corresponds to the majority of homozygous *k*-mers of the diploid (2n) genome, the smaller peak at half the coverage comprises haplotype-specific (1n) *k*-mers. Small peaks at 3n and 4n represent regions of higher copy numbers. Low-coverage *k*-mers derive from sequencing errors and bacterial contamination. Cumulatively, the *k*-mer distribution suggests an approximate haploid genome size of 40 Mbp.

We annotated 82-84% of universal eukaryotic single-copy marker genes in each genome (Fig. 3). The majority of the missing markers are consistently absent from all four genomes, suggesting poor representation in reference databases or the complete lack of these genes from the group rather than problems with the quality of the underlying assemblies as the most likely explanation. A maximum-likelihood phylogeny reconstructed from a concatenated alignment of 123 shared single-copy markers suggests that RCC970-E3 and Cflag diverged most recently (Fig. 1b). The exact placement of the other two strains within the group could only be determined with low bootstrap support.

**Figure 3.**
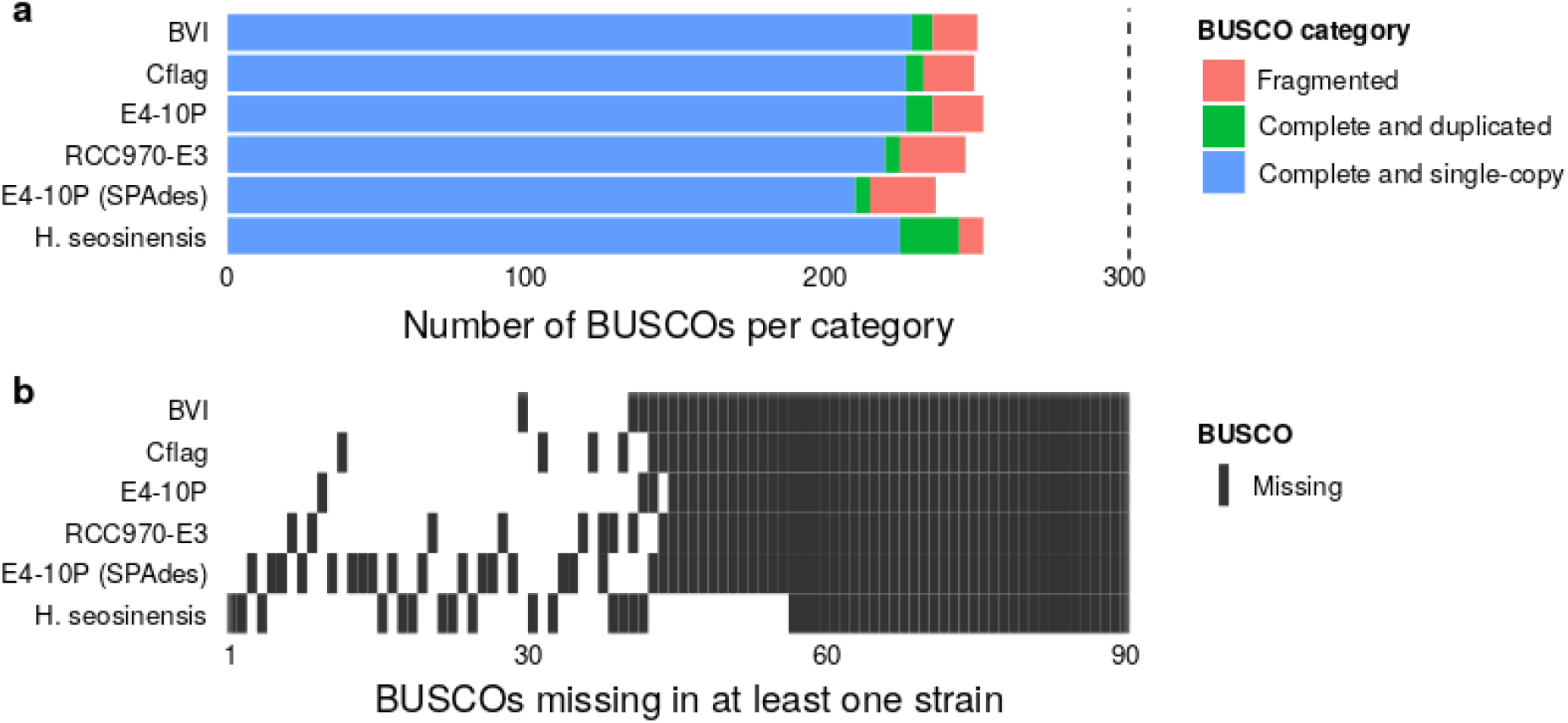
Completeness assessment for the genome assemblies based on single-copy orthologs. (**a**) Abundance of 303 single-copy core gene markers (BUSCOs) in different categories and assemblies. (**b**) Distribution of BUSCOs missing in at least one assembly (black tiles).

To analyze the genomic capabilities of *C. roenbergensis*, we annotated 7857-8483 protein-coding genes per strain using a combination of *de novo*, homology- and RNA-seq-supported gene prediction and homology-based functional assignments against UniProtKB/Swiss-Prot and the EggNOG database. In addition, all four assemblies comprise a curated circular mitochondrial genome that was annotated independently using a non-standard genetic code with UGA coding for tryptophan instead of a stop codon^13^.

We anticipate that the genomic data presented here will enable more detailed studies of *C. roenbergensis* shedding new light on this ecologically important group of marine grazers. Furthermore, the data may provide new insights into how asexually reproducing organisms maintain genome integrity and evolvability, and how this relates to their interactions with mobile genetic elements.

## Methods

### Strain maintenance, sample preparation, and sequencing

We selected and sequenced four strains of *C. roenbergensis* isolated from different locations in the Atlantic (Woods Hole, MA, USA; British Virgin Islands) and the Pacific (Yaquina Bay, OR, USA; South Pacific Ocean, 2200 km off the coast of Chile) (Table 1). All flagellate strains were grown in f/2 artificial seawater medium in the presence of either sterilized wheat grains (stock cultures) or 0.05% (w/v) yeast extract (for rapid growth), to stimulate the growth of a mixed bacterial community, which serve as a food source for *C. roenbergensis*. Each *C. roenbergensis* culture was subject to three consecutive rounds of single-cell dilution to obtain clonal strains as described previously^11^. For genome sequencing, suspension cultures of 2 L for each strain were grown to approximately 1×10^6^ cells/mL, then diluted two-fold with antibiotics-containing medium (30 µg/mL Streptomycin, 60 µg/mL Neomycin, 50 µg/mL Kanamycin, 50 µg/mL Ampicillin, 25 µg/mL Chloramphenicol) and incubated for 24 h at 22 °C and 60 rpm shaking to reduce the bacterial load. Cultures were filtered through a 100 μm Nitex mesh to remove large aggregates and centrifuged in various steps to further remove bacteria. First, the cultures were centrifuged for 40 min at 6000 x g and 20 °C (F9 rotor, Sorvall Lynx centrifuge), and the cell pellets were resuspended in 50 mL of f/2 medium, transferred to 50 mL polycarbonate tubes and centrifuged for 10 min at 4500 x g, 20 ◦C in an Eppendorf 5804R centrifuge. The supernatant was discarded and the cell pellet was resuspended in 50 mL of PBS (phosphate-buffered saline) medium. This washing procedure was repeated 10 times until the supernatant was clear, indicating that most bacteria had been removed. In the end, the flagellates were pelleted and resuspended in 2 mL of PBS medium. Genomic DNA from approximately 1×10^9^ cells of strains was isolated using the Blood & Cell Culture DNA Midi Kit (Qiagen, Hilden, Germany). The genomes were sequenced on an Illumina MiSeq platform (Illumina, San Diego, California, USA) using the MiSeq reagent kit version 3 at 2×300-bp read length configuration. The E4-10P genome was sequenced by GATC Biotech AG (Constance, Germany) with the standard MiSeq protocol. The E4-10M1, BVI, Cflag and RCC970-E3 genomes were prepared and sequenced at the Max Planck Genome Centre (Cologne, Germany) with NEBNext High-Fidelity 2×PCR Master Mix chemistry and a reduced number of enrichment PCR cycles (six) to reduce AT-bias. We also sequenced genomic DNA of all strains on a Pacific Biosciences RS II platform (2-3 SMRT cells each, Max Planck Genome Centre, Cologne, Germany).

### Assembly, decontamination, and refinement

MiSeq reads were trimmed for low-quality bases and adapter contamination using Trimmomatic^14^. PacBio reads were extracted from the raw data files with DEXTRACTOR^15^. Proovread^16^ was used for the hybrid correction of the PacBio reads with the respective trimmed MiSeq read sets. K-mer analysis was carried out with jellyfish^17^ and custom R scripts to plot and the distribution and estimate the genome size (Data Citation 3). To determine the best assembly strategy, we assessed draft assemblies generated with different approaches as described in the Technical Validation section. The improved high-quality drafts presented here were assembled using Canu v1.8^18^ from raw PacBio reads only for CrE4-10P, CrBVI, and CrCflag, and from raw and Illumina-corrected PacBio reads for CrRCC970-E3. For the latter strain, raw and corrected versions of the same PacBio reads were used together to mitigate low PacBio coverage in this particular sample and obtain a more contiguous assembly. Following the initial assembly, we used Redundans^19^ to remove redundant contigs, which were reconstructed as individual alleles due to high heterozygosity. To reduce misassemblies, we further broke up contigs at unexpected drops in coverage based on reads mapped with minimap2^20^ and identified with the custom Perl script bam-junctions (Data Citation 3) and bedtools^21^. After exploring different approaches (see Technical Validation) bacterial contamination was identified and removed based on taxonomic assignments generated with Kaiju^22^ and the script tax-resolve (Data Citation 3) using the ETE 3 python library(Huerta-Cepas et al. 2016). To obtain these assignments, each contig was split into 500 bp fragments, which we classified against the NCBI non-redundant protein database. Contigs with more than 50% of fragments annotated as bacteria were excluded from the assembly. Finally, to remove base-level errors we polished the assemblies in two rounds by mapping back first PacBio, then Illumina reads with minimap2^20^, and by generating consensus sequences from the mappings with Racon^23^.

### Gene prediction and functional annotation

Repetitive regions were detected with WindowMasker^24^. Only repetitive regions with a minimum length of 100 bp were retained. tRNA genes were predicted with tRNAscan2.0^25^ with a minimum score of 70. Gene prediction was performed with the BRAKER pipeline^26,27^, which utilizes BLAST ^28,29^, Augustus^30,31^ and GeneMark-ES^32,33^. Augustus and GeneMark-ES gene models were trained with publicly available transcriptomic data of *C. roenbergensis* E4-10P as extrinsic evidence (Data Citation 4)^34^. Prior to gene prediction, splice sites were detected by HISAT^35^ and processed with samtools^36,37^. Protein functions were assigned with two different approaches: 1) by blastx best hit against the UniProtKB/Swiss-Prot database v14/01/2019^38^ and 2) by eggNOG-mapper^39^ best match against the EggNOG v4.5.1 database^40^. Only results with an E-value of 10^−3^ or lower were retained. In addition, blastx hits with bitscores below 250, percentage identities below 30, or raw scores below 70 were ignored.

### Mitochondrial genome curation and annotation

To obtain a complete and correctly annotated mitochondrial genome, we first mapped the whole genome assemblies against an existing *C. roenbergensis* mitochondrial reference genome (NCBI accession NC_000946.1) to identify the mitochondrial contig using minimap2^20^. We then extracted the contig, trimmed overlapping ends of the circular sequence with seq-circ-trim and reset the start to the same location as the reference genome - the large subunit ribosomal RNA gene - with seq-circ-restart (Data Citation 3). Gene annotation was carried out with Prokka^41^ and with an adjusted non-standard translation code. Predicted tRNA and coding genes completely overlapping other coding regions were manually removed guided by the reference genome annotation.

### PCR and reverse-transcription PCR conditions

Genomic DNA (gDNA) was extracted from 200 μl of suspension culture with the QIAamp DNA Mini kit (Qiagen, Hilden, Germany) following the manufacturer’s instructions for DNA purification of total DNA from cultured cells, with a single elution step in 100 μl of double-distilled (dd) H_2_O and storage at −20 °C.

For extraction of total RNA, 500 μl of suspension culture were centrifuged for 5 min at 4,500*g*, 4 °C. The supernatants were discarded and the cell pellets were immediately flash-frozen in N_2_(l) and stored at −80 °C until further use. RNA extraction was performed with the Qiagen RNeasy Mini Kit following the protocol for purification of total RNA from animal cells using spin technology. Cells were disrupted with QIAshredder homogenizer spin columns and an on-column DNase I digest was performed with the Qiagen RNase-Free DNase Set. RNA was eluted in 50 μl of RNase-free molecular biology grade water. The RNA was then treated with 1 μl TURBO DNase (2 U/μl) for 30 min at 37 °C according to the manufacturer’s instructions (Ambion via ThermoFisher Scientific, Germany). RNA samples were analyzed for quantity and integrity with a Qubit 4 Fluorometer (Invitrogen via ThermoFisher Scientific, Germany) using the RNA Broad Range and RNA Integrity and Quality kit respectively.

For cDNA synthesis, 6 μl of each RNA sample was reverse transcribed using the Qiagen QuantiTect Reverse Transcription Kit according to the manufacturer’s instructions. This protocol included an additional DNase treatment step and the reverse transcription reaction using a mix of random hexamers and oligo(dT) primers. Control reactions to test for gDNA contamination were done for all samples by adding ddH_2_O instead of reverse transcriptase to the reaction mix. The cDNA was diluted fivefold with RNase-free H_2_O and analyzed by PCR with gene-specific primers (Table 4).

**Table 4.**
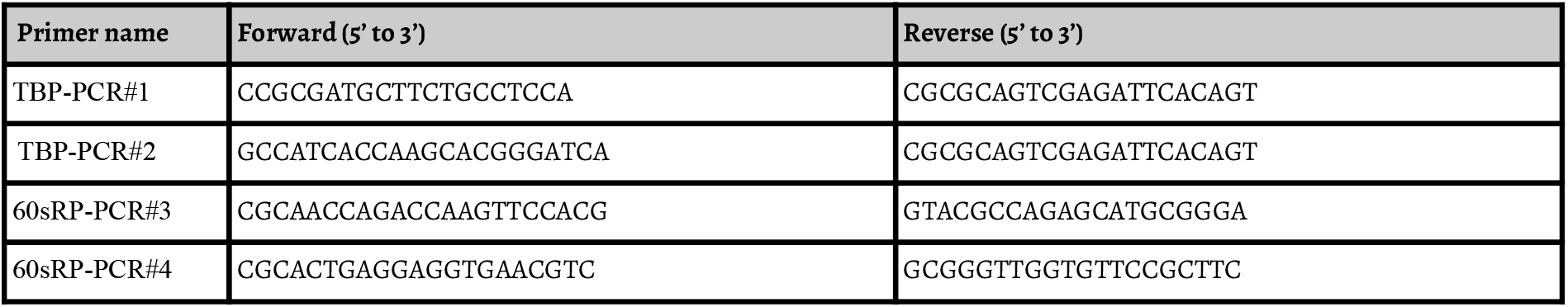
Primers used for the validation of the intron-exon structure in two genes of *C. roenbergensis* RCC_970_-E_3_.

PCR amplifications were performed using 2 ng of gDNA template or 2 μl of the diluted cDNA in a 20 μl reaction mix containing 10 μl Platinum™ II Hot-Start PCR Master Mix (Invitrogen via ThermoFisher Scientific, Germany), 4 μl Platinum GC Enhancer and 0.2 μM of each primer.

The PCRs were performed in a ProFlex PCR System (Applied Biosystems via ThermoFisher Scientific, Germany) with the following cycling conditions: 2 min denaturation at 94 °C and 35 cycles of 15 s denaturation at 94 °C, 20 s annealing at 60 °C (for all primers) and 20 s extension at 68 °C. For product analysis, 1 μl of each reaction were mixed with loading dye and pipetted on a 1% (w/v) agarose gel supplemented with GelRed. The marker lanes contained 0.5 μg of GeneRuler 100 bp DNA Ladder (Fermentas, Thermo-Fisher Scientific, USA). The gel was electrophoresed for 1 h at 100 V and visualized on a ChemiDoc MP Imaging System (BioRad, Germany).

### Phylogenetic analysis

Phylogenetic relationships among the *C. roenbergensis* strains and *Halocafeteria seosinensis* (Genbank accession LVLI00000000) were reconstructed from the concatenated alignment of 123 shared single-copy orthologous genes. Orthologs were identified with BUSCO^42,43^ with the eukaryotic lineage dataset. Only orthologs present in all five genomes with a BUSCO score of at least 125 and a minimum covering sequence length of 125 bp were taken into account. Orthologous protein sequences were aligned with MAFFT^44^ and trimmed for poorly aligned regions with trimAl^45^. The phylogenetic tree was computed with RAxML using the GAMMA model of rate heterogeneity and automatically determined amino acid substitution models for each partition. The bootstrap confidence values were computed with 100 iterations of rapid bootstrapping. The tree was rooted with *H. seosinensis* as the phylogenetic outgroup using phytools^46^ and visualized with ggtree^47^.

### Data Records

The raw Illumina and PacBio sequencing reads are available from the NCBI Sequence Read Archive (Data Citation 1). Accession numbers, library size, and coverage statistics can be found in Table 2. The curated and annotated assemblies for E4-10P, BVI, Cflag ad RCC970-E3 have been deposited as Whole Genome Shotgun projects at DDBJ/ENA/GenBank under the accessions VLTO00000000, VLTN00000000, VLTM00000000, VLTL00000000. The versions described in this paper are version VLTO01000000, VLTN01000000, VLTM01000000, VLTL01000000 (Data Citations 2). The draft assemblies we initially generated to determine the best assembly strategy are available from GitHub and Zenodo (Data Citation 3) together with custom code used in the analysis.

### Technical Validation

Overall sequencing quality of MiSeq and PacBio read data was assessed with FastQC v0.11.3^48^. To choose the best assemblies for further refinement, we evaluated different assemblers and alternative assembly strategies. In particular, we assembled draft genomes using MiSeq reads and corrected PacBio data with SPAdes^49^ in diploid mode, and generated assemblies from raw PacBio reads with Flye^50^ and wtdbg2^51^. Using QUAST^52^, BUSCO^43^, and the misassembly detection procedure described in Material and Methods, we assessed contiguity, completeness, and quality of the different assemblies. We found that the Illumina-based assemblies, in general, were less complete and less contiguous than the PacBio-based assemblies (for best SPAdes assembly see Data Citation 3). The PacBio assemblies differed primarily in the number of potential misassembly sites, with Canu being least prone to this potential issue. Therefore, we selected assemblies generated with Canu for further processing and analysis.

Because the read data was obtained from non-axenic cultures, we screened the assemblies carefully for contaminations with a custom R script. Initially, we considered four different criteria: tetra-nucleotide frequencies, coverage, GC-content, and taxonomic assignments (Supplementary Fig. 1-4). Tetra-nucleotide frequencies and GC-content were computed with seq-comp (Data Citation 3). Medium contig coverage was determined based on MiSeq reads mapped with minimap2 using bam-coverage (Data Citation 3). Taxonomic assignments were generated with Kaiju^22^ (See Materials and Methods for details). We found that all four criteria generated similar and consistent results. All identified contaminations were classified as bacterial. Viral signatures could all be attributed to endogenous viral elements expected to be present in *Cafeteria* genomes. From this analysis, we established a simple rule for decontamination of the assemblies: Contigs with more than 50% of annotated regions classified as bacteria were excluded.

To further assess the completeness and quality of our assemblies, we used BUSCO^42,43^ to detect universal eukaryotic orthologous genes (Fig 1). We found that all our PacBio-based assemblies contain 82%-84% of the expected 303 orthologs. Moreover, most missing orthologs are absent from all four assemblies suggesting poor representation in the database or the complete lack of some of these markers from this group as a systemic issue, rather than assembly problems, which would affect different genes in different assemblies. To validate the automated gene predictions, we spot-checked the intron-exon structure for two genes using regular and reverse-transcription PCR (Supplementary Fig. 5). We selected two intron-containing genes, namely genes coding for a TATA-binding protein (locus tag: FNF27_01237) and a 60S ribosomal protein (locus tag: FNF28_01226). Primers were designed to amplify the intronic regions and PCR with these primers resulted in long amplicons when using gDNA as the template and a shorter amplicon when using cDNA as the template. The cDNA was obtained after reverse transcription of total RNA from *C. roenbergensis* strain RCC970-E3.

### Code Availability

All custom code used to generate and analyze the data presented here is available from Data Citation 3 and from http://github.com/thackl/cr-genomes.

### Software versions and relevant parameters

Trimmomatic v0.32 (ILLUMINACLIP:TruSeq3-PE.fa:2:30:10 SLIDINGWINDOW:10:20 MINLEN:75 LEADING:3 TRAILING:3); proovread v2.12 (config settings: ‘seq-filter’ => {‘--trim-win’ =>‘10,1’, ‘--min-length’ => 500}, ‘sr-sampling’ => {DEF => 0}); Canu v1.8; Flye v2.3.7; WTDBG v2.1; SPAdes v3.6.1 (--diploid); minimap2 v2.13-r858-dirty (PacBio reads: -x map-pb; MiSeq readsL -x sr); bam-junctions SHA: 28dc943 (-a200 -b200 -c5 -d2 -f30 -e30); Redundans v0.14a (--noscaffolding --norearrangements --nogapclosing); Kaiju v1.6.3 (-t kaijudb/nodes.dmp -f kaijudb/kaiju_db_nr_euk.fmi); Prokka v1.13 (--kingdom Mitochondria --gcode 4); DEXTRACTOR rev-844cc20; jellyfish v2.2.4; samtools v1.7; Racon v1.3.1; BUSCO v3.1.0; WindowMasker 1.0.0; tRNAscan v2.0; BRAKER v.2.1.1; BLAST v.2.6.0+; Augustus v3.3.2; GeneMark-ES v.4.38; HISAT v.2.1.0; MAFFT v7.310; trimAl v1.4.rev22 (-strictplus); RAxML v8.2.9 (-p 13178 -f a -x 13178 -N 100 -m PROTGAMMAWAG -q part.txt); phytools v0.6-60; ggtree v1.14.4;

## Acknowledgments

This work was supported by the Max Planck Society and grants from the Gordon and Betty Moore Foundation (Grant ID: 5734), ASSEMBLE (Grant ID: 227799), and the European Regional Development Fund, EFRE-Program, European Territorial 190 Cooperation (ETZ) 2014-2020, Interreg V A, Project 41. We thank the staff of the Max Planck Genome Centre Cologne for excellent assistance with sequencing, Daniel Vaulot & the team of the Roscoff Culture Collection and Dave Caron for providing flagellate strains (RCC970 and Cflag, respectively), Ilme Schlichting and Curtis Suttle for providing samples (BVI and E4-10, respectively) and advice, Chris Roome for IT assistance, and Alexa Weinmann for technical support. All BLAST computations were performed on the MaRC2 high-performance cluster of the University of Marburg, which is supported by the State Ministry of Higher Education, Research and the Arts. We especially thank Mr. Sitt of HPC-Hessen for his technical support.

## Author contributions

T.H. and M.G.F. designed the study and wrote the manuscript with contributions from all other authors. K.B. and M.G.F. maintained the cultures and extracted DNA for sequencing. T.H. generated, curated and analyzed the assemblies and annotation data. R.M. carried out gene annotations and phylogenetic analyses. S.D. conducted the experimental validation of gene models. M.G.F and D.H. supervised the project.

## References

1. del Campo, J. et al. The others: our biased perspective of eukaryotic genomes. Trends Ecol. Evol. 29, 252–259 (2014).

2. Fenchel, T. & Patterson, D. J. Cafeteria roenbergensis nov. gen., nov. sp., a heterotrophic microflagellate from marine plankton. Mar. Microb. Food Webs (1988).

3. Larsen, J. & Patterson, D. J. Some flagellates (Protista) from tropical marine sediments. J. Nat. Hist. 24, 801–937 (1990).

4. Patterson, D. J., Nygaard, K., Steinberg, G. & Turley, C. M. Heterotrophic flagellates and other protists associated with oceanic detritus throughout the water column in the mid North Atlantic. J. Mar. Biol. Assoc. U. K. 73, 67–95 (1993).

5. Atkins, M. S., Teske, A. P. & Anderson, O. R. A survey of flagellate diversity at four deep-sea hydrothermal vents in the Eastern Pacific Ocean using structural and molecular approaches. J. Eukaryot. Microbiol. 47, 400–411 (2000).

6. de Vargas, C. et al. Eukaryotic plankton diversity in the sunlit ocean. Science 348, 1261605 (2015).

7. Harding, T., Brown, M. W., Simpson, A. G. B. & Roger, A. J. Osmoadaptative Strategy and Its Molecular Signature in Obligately Halophilic Heterotrophic Protists. Genome Biol. Evol. 8, 2241–2258 (2016).

8. Fischer, M. G. & Suttle, C. A. A virophage at the origin of large DNA transposons. Science 332, 231–234 (2011).

9. Fischer, M. G., Allen, M. J., Wilson, W. H. & Suttle, C. A. Giant virus with a remarkable complement of genes infects marine zooplankton. Proc. Natl. Acad. Sci. U. S. A. 107, 19508–19513 (2010).

10. Krupovic, M., Kuhn, J. H. & Fischer, M. G. A classification system for virophages and satellite viruses. Arch. Virol. (2015). doi:10.1007/s00705-015-2622-9

11. Fischer, M. G. & Hackl, T. Host genome integration and giant virus-induced reactivation of the virophage mavirus. Nature 540, 288–291 (2016).

12. Le Gall, F. et al. Picoplankton diversity in the South-East Pacific Ocean from cultures. (2008). doi:10.5194/bg-5-203-2008

13. Gray, M. W. et al. Genome structure and gene content in protist mitochondrial DNAs. Nucleic Acids Res. 26, 865–878 (1998).

14. Bolger, A. M., Lohse, M. & Usadel, B. Trimmomatic: a flexible trimmer for Illumina sequence data. Bioinformatics (2014).

15. Myers, G. Efficient local alignment discovery amongst noisy long reads. in Lecture Notes in Computer Science (including subseries Lecture Notes in Artificial Intelligence and Lecture Notes in Bioinformatics) 8701 LNBI, 52–67 (2014).

16. Hackl, T., Hedrich, R., Schultz, J. & Förster, F. proovread: large-scale high-accuracy PacBio correction through iterative short read consensus. Bioinformatics 30, 3004–3011 (2014).

17. Marçais, G. & Kingsford, C. A fast, lock-free approach for efficient parallel counting of occurrences of k-mers. Bioinformatics 27, 764–770 (2011).

18. Koren, S., Walenz, B. P., Berlin, K., Miller, J. R. & Phillippy, A. M. Canu: scalable and accurate long-read assembly via adaptive k-mer weighting and repeat separation. bioRxiv 071282 (2016). doi:10.1101/071282

19. Pryszcz, L. P. & Gabaldón, T. Redundans: an assembly pipeline for highly heterozygous genomes. Nucleic Acids Res. 44, e113 (2016).

20. Li, H. Minimap 2: pairwise alignment for nucleotide sequences. Bioinformatics 34, 3094–3100 (2018).

21. Quinlan, A. R. BEDTools: The Swiss-Army Tool for Genome Feature Analysis. Curr. Protoc. Bioinformatics 47, 11.12.1–34 (2014).

22. Menzel, P., Ng, K. L. & Krogh, A. Fast and sensitive taxonomic classification for metagenomics with Kaiju. Nat. Commun. 7, 11257 (2016).

23. Vaser, R., Sović, I., Nagarajan, N. & Šikić, M. Fast and accurate de novo genome assembly from long uncorrected reads. Genome Res. 27, 737–746 (2017).

24. Morgulis, A., Gertz, E. M., Schäffer, A. A. & Agarwala, R. WindowMasker: window-based masker for sequenced genomes. Bioinformatics 22, 134–141 (2006).

25. Lowe, T. M. & Chan, P. P. tRNAscan-SE On-line: integrating search and context for analysis of transfer RNA genes. Nucleic Acids Res. 44, W54–7 (2016).

26. Lomsadze, A., Burns, P. D. & Borodovsky, M. Integration of mapped RNA-Seq reads into automatic training of eukaryotic gene finding algorithm. Nucleic Acids Res. 42, e119 (2014).

27. Hoff, K. J., Lange, S., Lomsadze, A., Borodovsky, M. & Stanke, M. BRAKER1: Unsupervised RNA-Seq-Based Genome Annotation with GeneMark-ET and AUGUSTUS. Bioinformatics 32, 767–769 (2016).

28. Altschul, S. F., Gish, W., Miller, W., Myers, E. W. & Lipman, D. J. Basic local alignment search tool. J. Mol. Biol. 215, 403–410 (1990).

29. Camacho, C. et al. BLAST+: architecture and applications. BMC Bioinformatics 10, 421 (2009).

30. Stanke, M., Schöffmann, O., Morgenstern, B. & Waack, S. Gene prediction in eukaryotes with a generalized hidden Markov model that uses hints from external sources. BMC Bioinformatics 7, 62 (2006).

31. Stanke, M., Diekhans, M., Baertsch, R. & Haussler, D. Using native and syntenically mapped cDNA alignments to improve de novo gene finding. Bioinformatics 24, 637–644 (2008).

32. Ter-Hovhannisyan, V., Lomsadze, A., Chernoff, Y. O. & Borodovsky, M. Gene prediction in novel fungal genomes using an ab initio algorithm with unsupervised training. Genome Res. 18, 1979–1990 (2008).

33. Lomsadze, A., Ter-Hovhannisyan, V., Chernoff, Y. O. & Borodovsky, M. Gene identification in novel eukaryotic genomes by self-training algorithm. Nucleic Acids Res. 33, 6494–6506 (2005).

34. Keeling, P. J. et al. The Marine Microbial Eukaryote Transcriptome Sequencing Project (MMETSP): illuminating the functional diversity of eukaryotic life in the oceans through transcriptome sequencing. PLoS Biol. 12, e1001889 (2014).

35. Kim, D., Langmead, B. & Salzberg, S. L. HISAT: a fast spliced aligner with low memory requirements. Nat. Methods 12, 357–360 (2015).

36. Li, H. et al. The Sequence Alignment/Map format and SAMtools. Bioinformatics 25, 2078–2079 (2009).

37. Barnett, D. W., Garrison, E. K., Quinlan, A. R., Strömberg, M. P. & Marth, G. T. BamTools: a C++ API and toolkit for analyzing and managing BAM files. Bioinformatics 27, 1691–1692 (2011).

38. UniProt Consortium. UniProt: a worldwide hub of protein knowledge. Nucleic Acids Res. 47, D506–D515 (2019).

39. Huerta-Cepas, J. et al. Fast Genome-Wide Functional Annotation through Orthology Assignment by eggNOG-Mapper. Mol. Biol. Evol. 34, 2115–2122 (2017).

40. Huerta-Cepas, J. et al. eggNOG 4.5: a hierarchical orthology framework with improved functional annotations for eukaryotic, prokaryotic and viral sequences. Nucleic Acids Res. 44, D286–93 (2016).

41. Seemann, T. Prokka: Rapid prokaryotic genome annotation. Bioinformatics 30, 2068–2069 (2014).

42. Simão, F. A., Waterhouse, R. M., Ioannidis, P., Kriventseva, E. V. & Zdobnov, E. M. BUSCO: assessing genome assembly and annotation completeness with single-copy orthologs. Bioinformatics 31, 3210–3212 (2015).

43. Waterhouse, R. M. et al. BUSCO applications from quality assessments to gene prediction and phylogenomics. Mol. Biol. Evol. (2017). doi:10.1093/molbev/msx319

44. Katoh, K. & Standley, D. M. MAFFT multiple sequence alignment software version 7: improvements in performance and usability. Mol. Biol. Evol. 30, 772–780 (2013).

45. Capella-Gutiérrez, S., Silla-Martínez, J. M. & Gabaldón, T. trimAl: a tool for automated alignment trimming in large-scale phylogenetic analyses. Bioinformatics 25, 1972–1973 (2009).

46. Revell, L. J. phytools: an R package for phylogenetic comparative biology (and other things). Methods Ecol. Evol. 3, 217–223 (2012).

47. Yu, G., Smith, D. K., Zhu, H., Guan, Y. & Lam, T. T.-Y. ggtree: an r package for visualization and annotation of phylogenetic trees with their covariates and other associated data. Methods Ecol. Evol. 8, 28–36 (2017).

48. Andrews, S. FastQC: A quality control tool for high throughput sequence data. (2010). Available at: http://www.bioinformatics.babraham.ac.uk/projects/fastqc/.

49. Bankevich, A. & Nurk, S. SPAdes: a new genome assembly algorithm and its applications to single-cell sequencing. Journal of … (2012).

50. Kolmogorov, M., Yuan, J., Lin, Y. & Pevzner, P. A. Assembly of long, error-prone reads using repeat graphs. Nat. Biotechnol. 37, 540–546 (2019).

51. Ruan, J. & Li, H. Fast and accurate long-read assembly with wtdbg2. bioRxiv 530972 (2019). doi:10.1101/530972

52. Gurevich, A., Saveliev, V., Vyahhi, N. & Tesler, G. QUAST: quality assessment tool for genome assemblies. Bioinformatics 29, 1072–1075 (2013).

## Data Citations

1. NCBI Sequence Read Archive SRP215872 (2019).

2. Hackl, T., Martin, R., Fischer M. G. Genomes of Cafeteria roenbergensis, strains E4-10P, BVI, Cflag, and RCC970-E3. GenBank VLTO01000000, VLTN01000000, VLTM01000000, VLTL01000000 (2019)

3. Hackl, T., Martin, R., Fischer M. G. Supplementary code and data for genomes of C. roenbergensis: E4-10P, BVI, Cflag and RCC970-E3 doi:10.5281/zenodo.3381417 (2019)

4. Cyverse Data Commons MMETSP0942 (2014)

