## Supplemental Figures 1-5 for "Four high-quality draft genome assemblies of the marine heterotrophic nanoflagellate *Cafeteria roenbergensis*"

For custom code and supplementary raw data see doi:10.5281/zenodo.3376248.

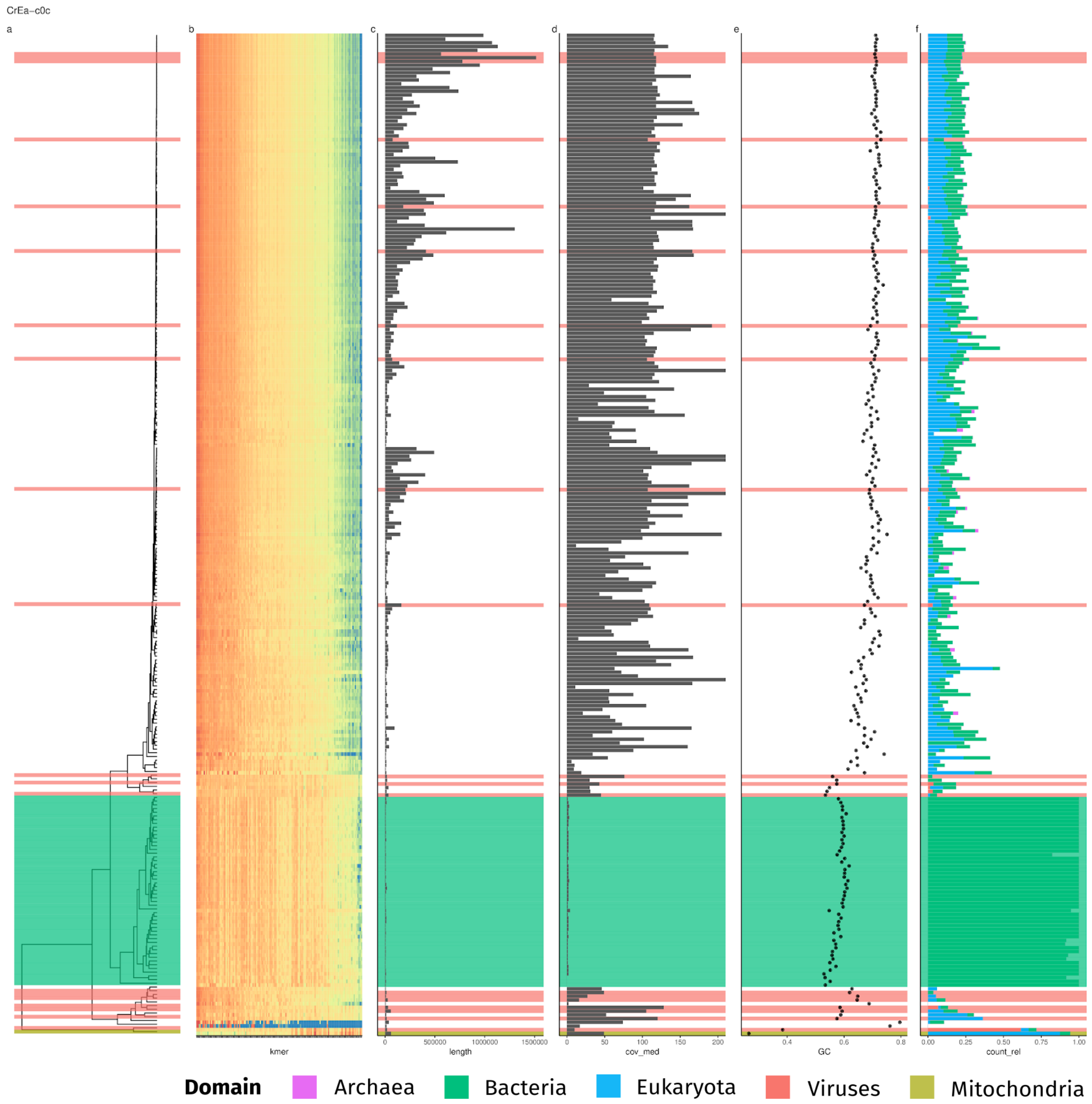


**Supplementary Figure 1. Contamination screening of assembly CrE410P.** (**a**) hierarchical clustering of contigs based on (**b**) scaled contig tetranucleotide frequencies. (**c**) Contig length. (**d**) Contig median coverage based on mapped Miseq reads. (**e**) Contig GC-content. (**f**) Distribution of taxonomic assignments at the domain level for 500 bp contig fragments. Colored bars present across all panels but (b) indicate sequences flagged as bacterial contamination (green), provirophage-containing (red) or mitochondrial (yellow).

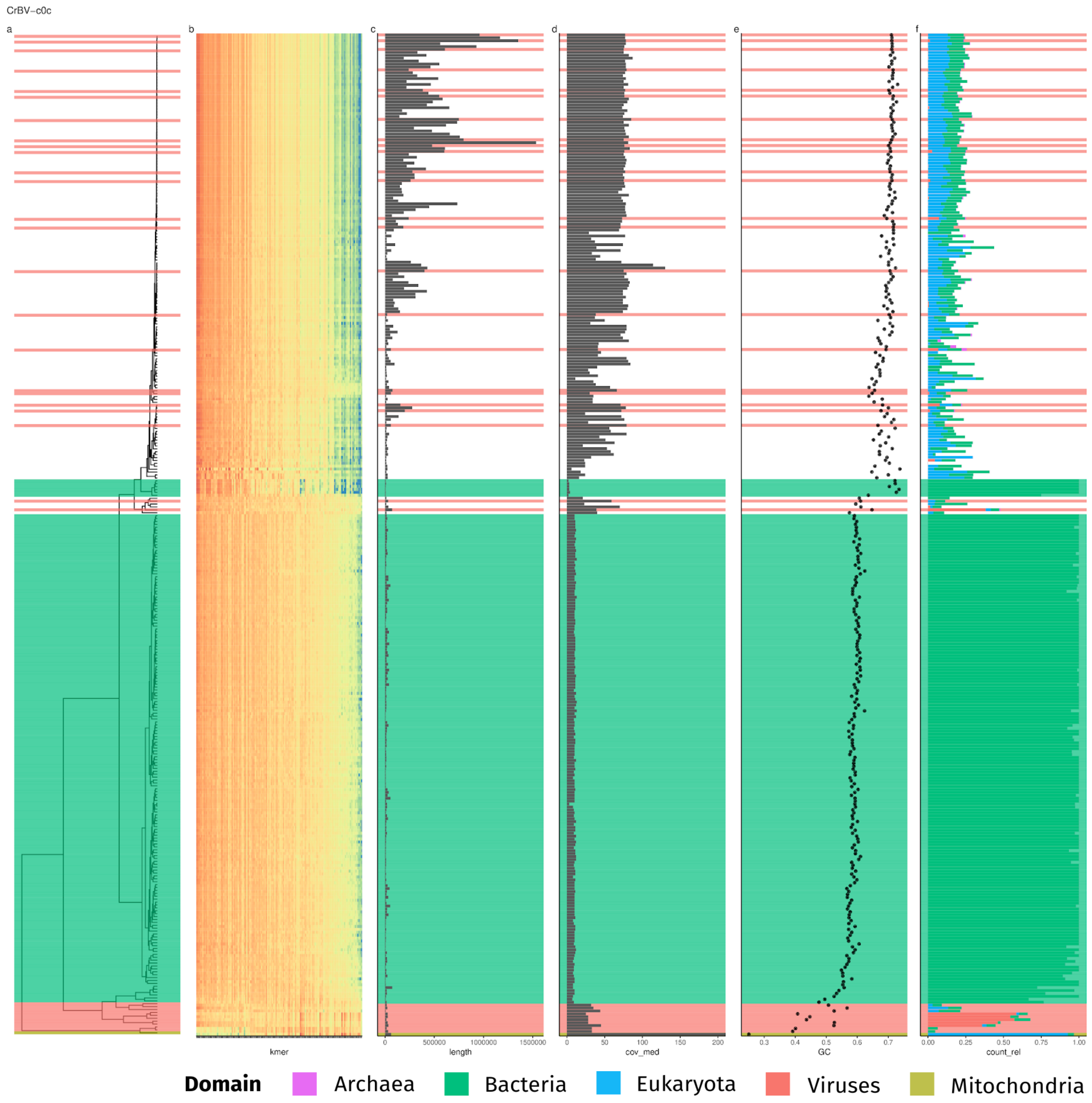


**Supplementary Figure 2. Contamination screening of assembly CrBVI.** (**a**) hierarchical clustering of contigs based on (**b**) scaled contig tetranucleotide frequencies. (**c**) Contig length. (**d**) Contig median coverage based on mapped Miseq reads. (**e**) Contig GC-content. (**f**) Distribution of taxonomic assignments at the domain level for 500 bp contig fragments. Colored bars present across all panels but (b) indicate sequences flagged as bacterial contamination (green), provirophage-containing (red) or mitochondrial (yellow).

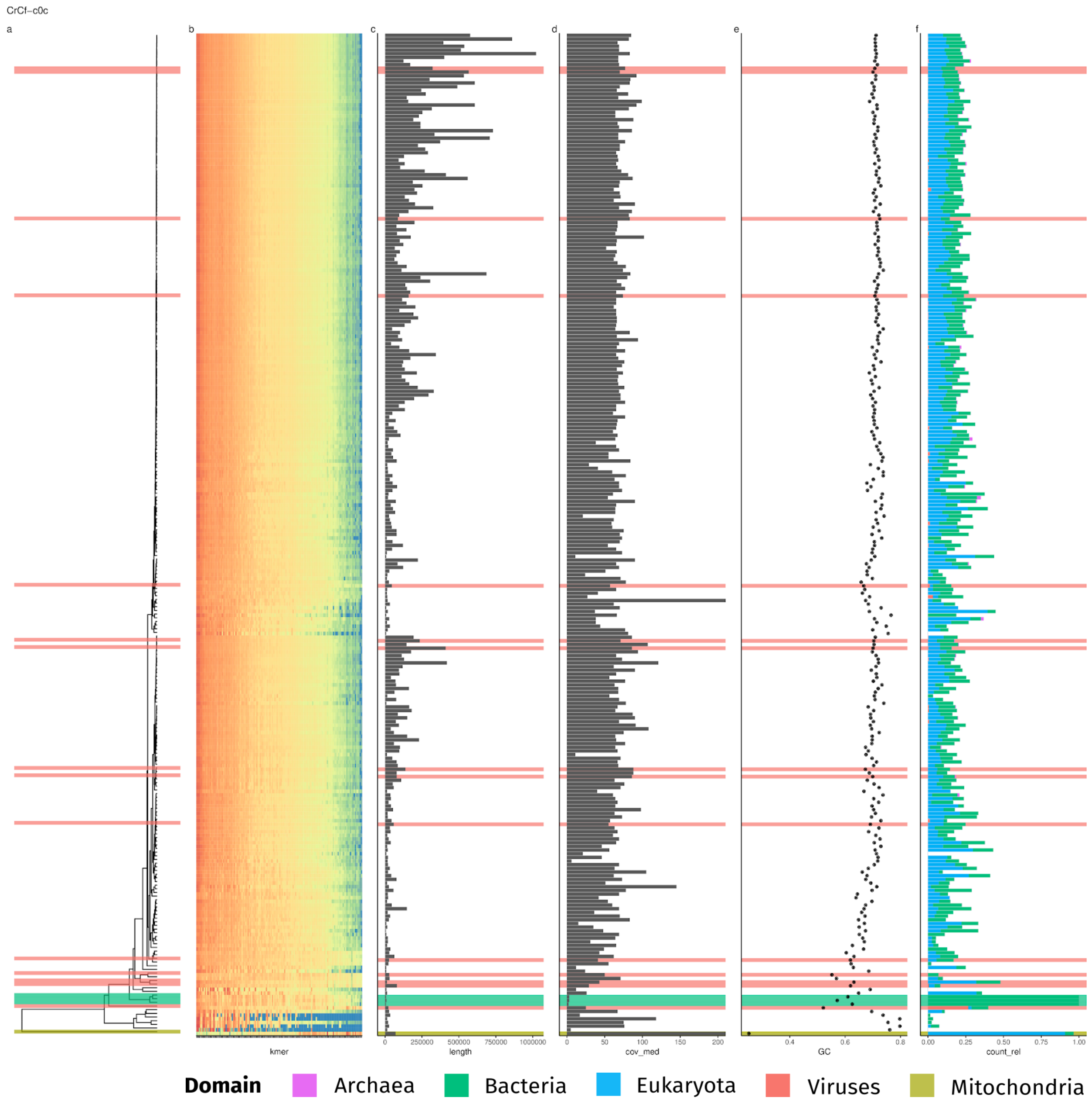


**Supplementary Figure 3. Contamination screening of assembly CrCflag.** (**a**) hierarchical clustering of contigs based on (**b**) scaled contig tetranucleotide frequencies. (**c**) Contig length. (**d**) Contig median coverage based on mapped Miseq reads. (**e**) Contig GC-content. (**f**) Distribution of taxonomic assignments at the domain level for 500 bp contig fragments. Colored bars present across all panels but (b) indicate sequences flagged as bacterial contamination (green), provirophage-containing (red) or mitochondrial (yellow).

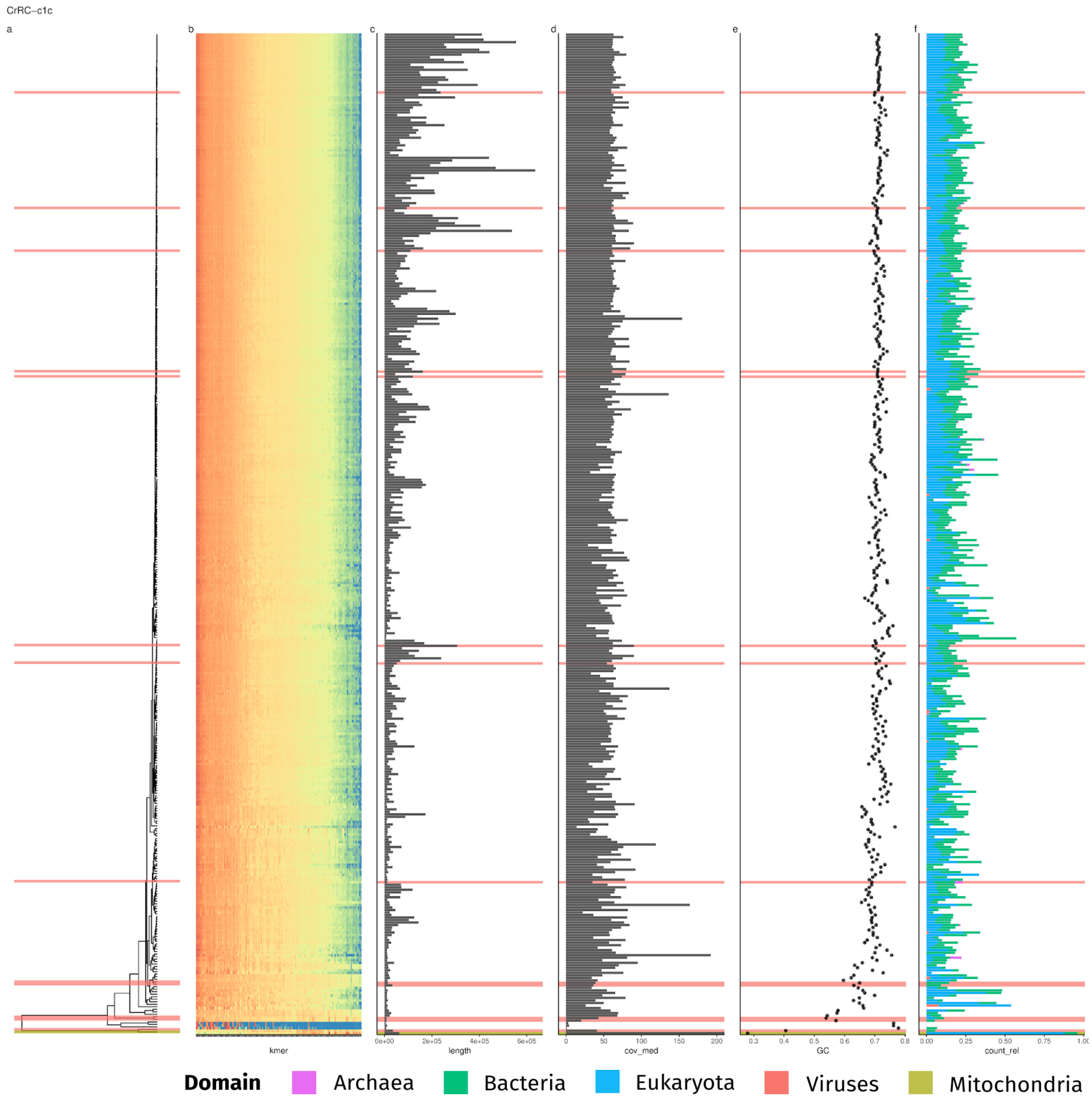


**Supplementary Figure 4. Contamination screening of assembly CrRCC970.** (**a**) hierarchical clustering of contigs based on (**b**) scaled contig tetranucleotide frequencies. (**c**) Contig length. (**d**) Contig median coverage based on mapped Miseq reads. (**e**) Contig GC-content. (**f**) Distribution of taxonomic assignments at the domain level for 500 bp contig fragments. Colored bars present across all panels but (b) indicate sequences flagged as bacterial contamination (green), provirophage-containing (red) or mitochondrial (yellow).

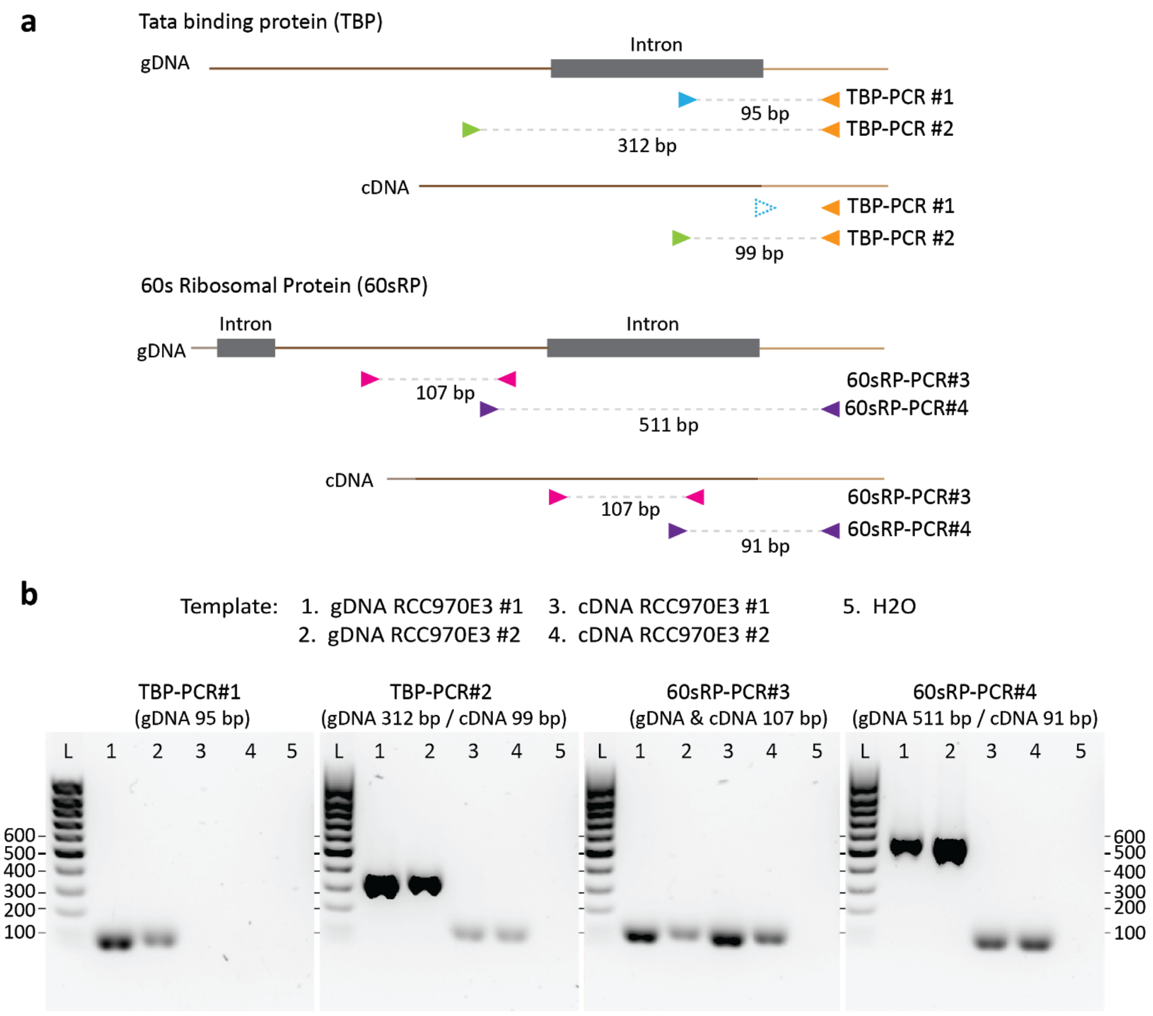


**Supplementary Figure 5. Intron validation for the TATA-binding protein and 60S ribosomal protein genes by PCR and reverse-transcription PCR.** (**a**) PCR primer design to amplify the intronic region. (**b**) Gel images of the PCR products obtained from gDNA or cDNA of *C. roenbergensis* strain RCC970-E3. Each condition was analyzed in biological duplicates. The lanes are labeled according to the templates listed above. L, DNA size standard.
